# A Structurally-Validated Multiple Sequence Alignment of 497 Human Protein Kinase Domains

**DOI:** 10.1101/776740

**Authors:** Vivek Modi, Roland L. Dunbrack

## Abstract

Studies on the structures and functions of individual kinases have been used to understand the biological properties of other kinases that do not yet have experimental structures. The key factor in accurate inference by homology is an accurate sequence alignment. We present a parsimonious, structure-based multiple sequence alignment (MSA) of 497 human protein kinase domains excluding atypical kinases, even those with related but somewhat different folds. The alignment is arranged in 17 blocks of conserved regions and unaligned blocks in between that contain insertions of varying lengths present in only a subset of kinases. The aligned blocks contain well-conserved elements of secondary structure and well-known functional motifs, such as the DFG and HRD motifs. From pairwise, all-against-all alignment of 272 human kinase structures, we estimate the accuracy of our MSA to be 97%. The remaining inaccuracy comes from a few structures with shifted elements of secondary structure, and from the boundaries of aligned and unaligned regions, where compromises need to be made to encompass the majority of kinases. A new phylogeny of the protein kinase domains in the human genome based on our alignment indicates that ten kinases previously labeled as “OTHER” can be confidently placed into the CAMK group. These kinases comprise the Aurora kinases, Polo kinases, and calcium/calmodulin-dependent kinase kinases.

## Introduction

Protein kinases catalyze the transfer of a phosphoryl group from an ATP molecule to substrate proteins^1^, and are crucial for cellular signaling pathways^2^. Mutations in kinases that lead to gain of function are frequently observed in many cancer types^3,4^, while mutations may also result in drug resistance rendering existing drugs inefficient^3^. Humans have over 500 genes that catalyze the phosphorylation of proteins, collectively called the ‘kinome’^5^.

Protein kinase activity is found in a number of protein families and superfamilies in the human proteome. The vast majority of human kinases come from one very large, diverse family that share a common fold consisting of an N-terminal lobe, composed of five β-sheet strands and an α-helix called the C-helix, and a C-terminal lobe comprising six α-helices^6^. The active site is located between the two lobes where the activation and catalytic loops form the ATP and substrate binding sites.

In 2002, Manning and coworkers identified 518 kinase genes in the human kinome^5^ which they divided into 478 typical kinase genes (13 of them containing two kinase domains, for a total of 491) and 40 atypical kinase genes. The annotation of the human genome has improved since the Manning paper; currently Uniprot identifies 483 human proteins containing 496 typical kinase domains (https://www.uniprot.org/docs/pkinfam). Uniprot identifies 29 atypical human kinases. Some of these are distantly related to the typical kinase domain, thus making them a superfamily, including Alpha kinases^7^, ADCK kinases^8^, RIO kinases^9^, FAM20C kinases^10^, and the PI3-PI4 kinase family, which contains the protein kinases ATM, ATR, and MTOR^11^. In addition, there are proteins that do not appear to share an evolutionary relationship with typical kinases that also phosphorylate proteins, such as pyruvate dehydrogenase kinases^12^.

For any large protein family, an accurate multiple sequence alignment is the basis of an accurate phylogeny^13^ and structural and functional inferences^14^. In 2002, Manning et al. built a phylogenetic tree of 491 typical kinase domains from a multiple sequence alignment created without using any kind of structural information^5^. The accompanying poster and image of this tree is still widely used in scientific papers and presentations on kinases^15^. Multiple sequence alignments of kinases have been used to extend structural and functional information from the kinases with known structures to those without known structures. This includes the conformations of active and inactive kinases^16-18^, predictions of substrate specificity^19^, and analysis of kinase-drug interactions^20,21^.

A common problem in multiple sequence alignments of large, diverse protein families is that they are very ‘gappy,’ i.e., containing many gap characters in every sequence in order to align inserted segments of different lengths that may be present in only a small subset of the sequences. This is true of the alignment used to produce the kinome tree of Manning et al.^5^ The gappy regions are usually present between major elements of secondary structure, where the family members may have widely divergent sequence loop lengths due to numerous insertions that occurred in different lineages of family members. The gappiness makes alignments difficult to visualize and produces errors in phylogenetic inference^22^.

In this paper, we present a parsimonious, structure-based multiple sequence alignment (MSA) of 497 human typical kinase domains from 484 human genes. We developed the MSA from pairwise structure alignment to a reference kinase (Aurora A kinase) and sequence alignment for kinases of unknown structure to their closest homologues of known structure. One of our central goals was to develop a parsimonious alignment of kinases containing as few gap regions as possible. Our alignment therefore contains aligned blocks (in upper case letters) that represent common structural elements, usually of secondary structure elements and important motifs, present in most or all of the kinases. These regions are separated by left-justified, unaligned sequence regions in lower case letters containing insertions of different lengths present in only subsets of the kinases. Our alignment contains 17 aligned blocks and 16 unaligned regions between them.

We validated this alignment with an all-against-all pairwise structure alignment of 272 protein kinases, and we show that the alignment is significantly more accurate than the Manning alignment and other recent alignments for classifying kinases^16,23^. Finally, we used our alignment to produce an accurate phylogeny of the kinase domains. Guided by the phylogeny and HMMs for each group we assign ten kinases previously categorized as “OTHER” by Manning et al. to the CAMK group, consisting of Aurora kinases, Polo-like kinases, and calcium/calmodulin-dependent kinase kinases.

## RESULTS

### Typical and atypical kinases

Kinase sequences were identified from the list of human kinases and pseudokinases provided by Uniprot^24^ (https://www.uniprot.org/docs/pkinfam), which divides ‘typical’ kinases into nine groups (AGC, CAMK, CK1, CMGC, NEK, RGC, STE, TKL, TYR) and a group of OTHER kinases. It also contains a list of ‘atypical’ kinases divided into six families (ADCK, Alpha-type, PI3-PI4-related, RIO, PDK/BCKDK, and FASTK). To identify any kinases which are not included in the Uniprot list, we searched all human Uniprot sequences with PSI-BLAST and each human kinase domain as query. As a result, we were able to identify Uniprot entry PEAK3_HUMAN as an additional human kinase. It was identified in searches starting with PRAG1 and PEAK1. The relationship with protein kinases was confirmed with hhpred^25^.

Manning et al. did not include any atypical kinase in their sequence alignment and phylogenetic tree. However, before discarding them from our dataset we wanted to examine the atypical kinase structures available from the PDB. Since Aurora A is a good representative of typical protein kinase domains, we have used its structure as a reference to identify structural similarities and differences between typical kinases and atypical kinases. The kinase domain of Aurora A has 251 amino acid residues with eight helices, seven β strands, and all the known conserved motifs without any unusually long insertions (Figure 1).

**Figure 1:**
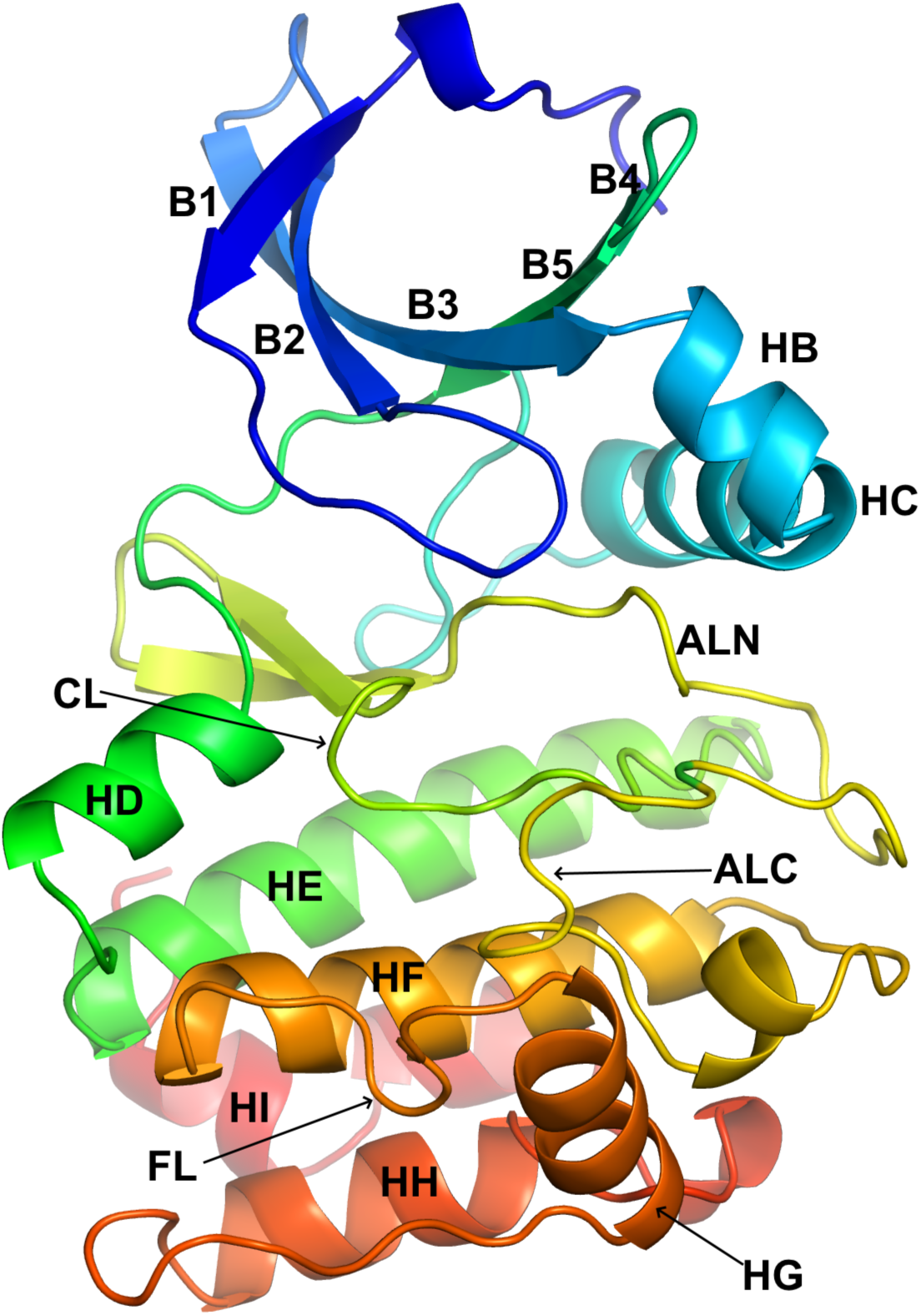
Structure of Aurora A (PDB: 3E5A_A) representing typical protein kinase domain. For the description of the labeled structural elements refer to Table 2.

Structure alignment with Aurora A indicated that four of the atypical kinase families are homologous to typical kinases (Figure 2), containing some elements of the typical kinase fold but containing changes and additions in elements of secondary structure. These include ADCK, Alpha-type, PI3-PI4-related, and RIO kinases. The ADCK (aarF-domain containing) kinases consist of five proteins: ADCK1, ADCK2, COQ8A (ADCK3), COQ8B (ADCK4), and ADCK5. Only the structure of COQ8A is available (PDB:4PED^8^). The structure consists of 384 residues, 13 helices, and eight β sheet strands. Structure alignment with FATCAT^26^ aligned 192 residues with an RMSD of 3.92 Å, covering the N-terminal domain, the HRD and DFG motifs, and the E and F helices of the C-terminal domain. COQ8A’s N-terminal domain contains an additional subdomain of five alpha helices, three of which precede the typical kinase domain and two of which are inserted between beta strand B3 and the C-helix. Instead of the activation loop leading into the F-helix, the DFG motif leads into a bundle of four alpha helices that precede COQ8A’s F-helix, which is followed by one additional helix.

**Figure 2:**
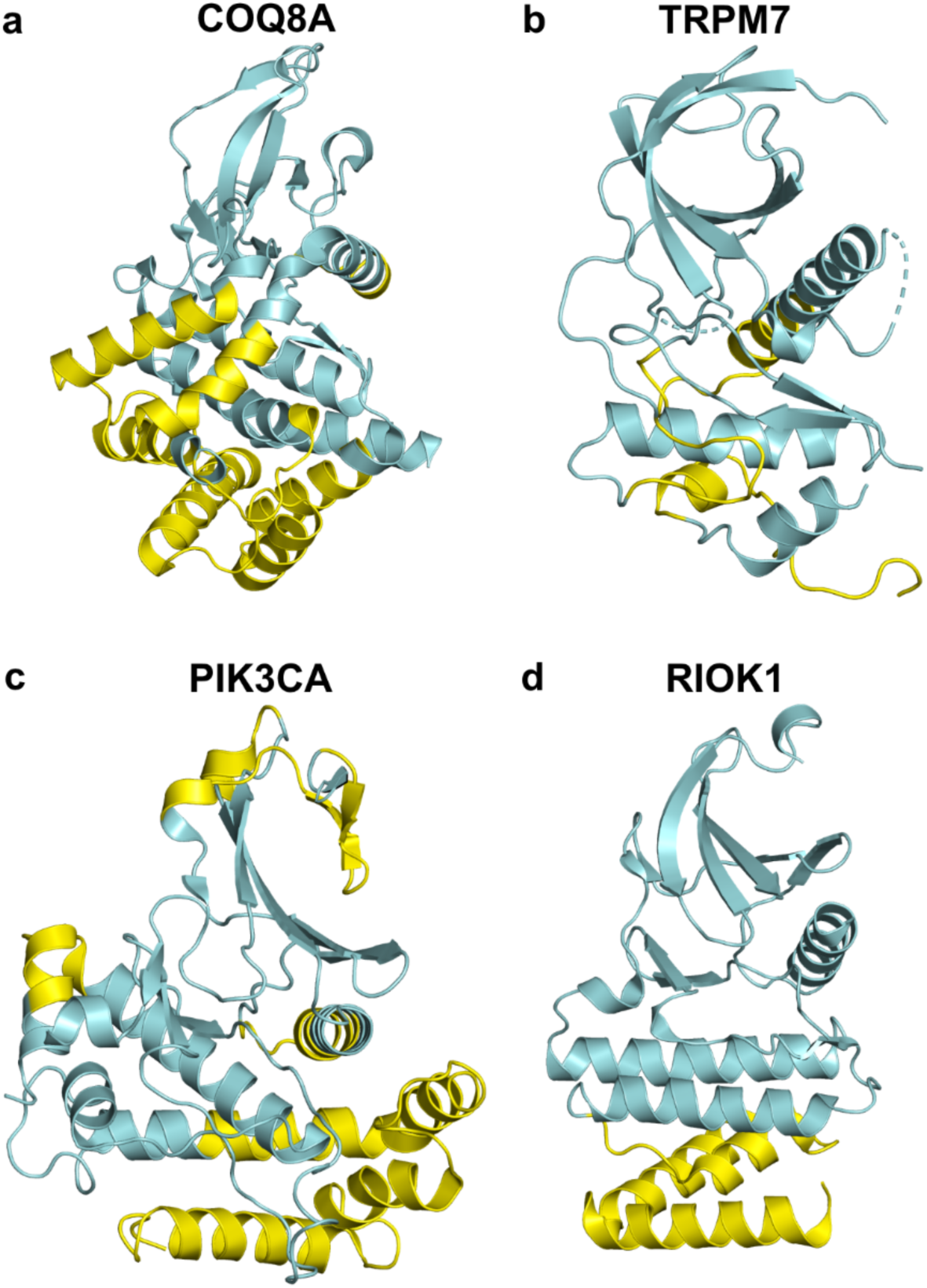
Representative structures from four different families of atypical kinases showing differences from typical kinase domains. **a)** ADCK protein kinase family - ADCK3 (4PED_A); **b)** Alpha-type - TRPM7 (1IAJ_A); **c)** PI3/PI4 - PIK3CA (4L2Y_A) and **d)** RIO-type Ser/Thr kinase family - RIOK1 (4OTP_A). The regions without any structural similarity to the typical kinase domain are colored in yellow, as identified by FATCAT and CEalign (in Pymol) after aligning to Aurora A (3E5A_A).

There are six kinases in the Alpha-type kinase family: ALPK1, ALPK2, ALPK3, EEF2K, TRPM6, and TRPM7. The structure of mouse TRPM7 (PDB:1IAJ^27^) has been determined; only the N-terminal domain and the E helix could be aligned to AURKA with an RMSD of 5.8 Å over 120 residues. The remainder of the C-terminal domain of TRMP7 consists of two beta sheet strands, large coil regions, and a short helix. The human PI3/PI4 kinases consist of seven genes: ATM, ATR, MTOR, PIK3CA, PIK3CB, PIK3CD, PIK3CG, PRKDC, and SMG1. All of these except SMG1 have known structures (PIK3CB is represented by a structure of mouse PIK3CB). The structure of PIK3CA (PDB: 4L2Y^28^) aligns with Aurora A with RMSD 6.0 Å over 168 residues. The structures of two of the three RIO kinases (RIOK1, RIOK2, and RIOK3) are known. The structures of RIOK1 (PDB: 4OTP^29^) and RIOK2 (PDB: 6FDN^30^) can be aligned with AURKA with 5.1 and 4.6 Å RMSD over 176 amino acids, because the orientation of the HG, HH, and HI helices are entirely different.

Two of the atypical protein kinase families listed by Uniprot do not appear to be homologous to typical kinases. The PDK/PCKDK protein kinase family consists of BCKDK, PDK1, PDK2, PDK3, and PDK4. The structures of all of these proteins are known (with BCKDK represented by rat BCKDK in the PDB by entry 3TZ5^31^ and PDK1 by 2Q8H^32^), and none of them resemble typical protein kinases. They consist of an N-terminal domain in the form of a bundle of four long α helices and a C-terminal domain of a five-stranded β sheet and three α helices. ECOD (Evolutionary Classification of Protein Domains) also does not classify these structures in the same homology group as the typical kinases^33^. While there is no structure of FAST kinase (Uniprot FASTK_HUMAN), the program hhpred^25^ found that the closest homologues in the PDB are restriction endonucleases (e.g., PDB:1CW0^34^), which do not appear to be homologous to typical kinases.

Overall, our examination indicated that every atypical family has significant differences from the typical kinase domain in the arrangement or presence of secondary structural elements. These differences make any alignment with the sequences of the typical kinase domain approximate and partial. Therefore, we did not include any atypical kinase sequence in our multiple sequence alignment of human protein kinases.

A summary of the 497 typical kinase domains from 484 Uniprot sequences included in our dataset and the available structures in the PDB are provided in Table 1. Thirteen kinases have two kinase domains in the sequence (see caption to Table 1), and some kinases were reassigned to different groups than the Uniprot or Manning designations (discussed below). The domain boundaries were initially identified using the domain annotations in Uniprot, while some of them were updated during the process of alignment. We labeled sequences by their HGNC gene names^35^ with a group name identifier from Uniprot. For example, the sequence for the gene AKT1 is labeled as AGC_AKT1 and EGFR is labeled TYR_EGFR. Some of the common kinases have gene names which are not easily recognized; for example VEGFR1, VEGFR2, and VEGFR3 are labeled TYR_FLT1, TYR_KDR, and TYR_FLT4 respectively. A table of Uniprot accession ids, Uniprot entry names, HGNC gene names, HGNC accessions, gene name synonyms, kinase group names, and domain boundaries is provided in Table S1. These sequences were used to create the MSA utilizing a variety of alignment programs and structural information.

**Table 1:** Number of kinase domains in each family in the multiple sequence alignment

|  | AGC | CAMK | CK1 | CMGC | NEK | RGC | STE | TKL | TYR | OTHER | Total |
| --- | --- | --- | --- | --- | --- | --- | --- | --- | --- | --- | --- |
| **Uniprot** | 62 | 88 | 12 | 65 | 11 | 5 | 47 | 42 | 90 | 66 | 484* |
| **Domains** | 62 | 92 | 12 | 65 | 11 | 5 | 47 | 42 | 94 | 67 | 497 |
| **Domains with PDB** | 29 | 49 | 10 | 39 | 3 | 0 | 27 | 25 | 64 | 28 | 274 |
| **Included in validation** | 29 | 49 | 10 | 39 | 3 | 0 | 27 | 25 | 64 | 26 | 272 |

\* 6 kinase genes have one AGC and one CAMK domain each (RPS6KA1, RPS6KA2, RPS6KA3, RPS6KA4, RPS6KA5 and RPS6KA6), and are counted twice in this table; 2 kinases have 2 CAMK domains (OBSCN, SPEG); 4 kinases have 2 TYR domains (JAK1, JAK2, JAK3, TYK2); 1 kinase has 2 domain in the OTHER category (EIF2AK4) ( $484+6+2+4+1=497$ domains). Two kinases AGC\_PDPK2P and AGC\_PRKY could be pseudogenes according to their Uniprot annotations.

### Forming a multiple sequence alignment of 497 human protein kinase domains

To examine the boundaries of conserved segments and common insertions across kinases, we first aligned the structure of Aurora A kinase N- and C-terminal domains separately to one representative structure of each of 271 other human kinases in the PDB using the program FATCAT^26^. We then used the SE program^36^ to read the structural alignments and print the corresponding sequence alignments. SE prints regions of low structural similarity (usually because of insertions or deletions) in lower case letters, and left-justifies them. An example is shown in Figure 3. To identify regions of structural variability in kinases, we counted the number of times a residue in the Aurora A sequence was structurally aligned across all the pairwise alignments output by SE (i.e. printed in upper case). This provides a count of how often each residue is structurally conserved, indicating the locations of insertions, deletions, and structural variations across the kinases (Figure 4). The tallest bars in the plot display the conserved regions in the alignments while the shorter bars represent low similarity regions or segments abutting common insertions and deletions. The region between ALN and ALC have shorter bars because the activation loop adopts very different conformations across kinase structures, resulting in relatively poor pairwise structural alignment in this region. We used these blocks of conserved regions and intermittent low similarity regions to guide the formation of the MSA.

**Figure 3:**
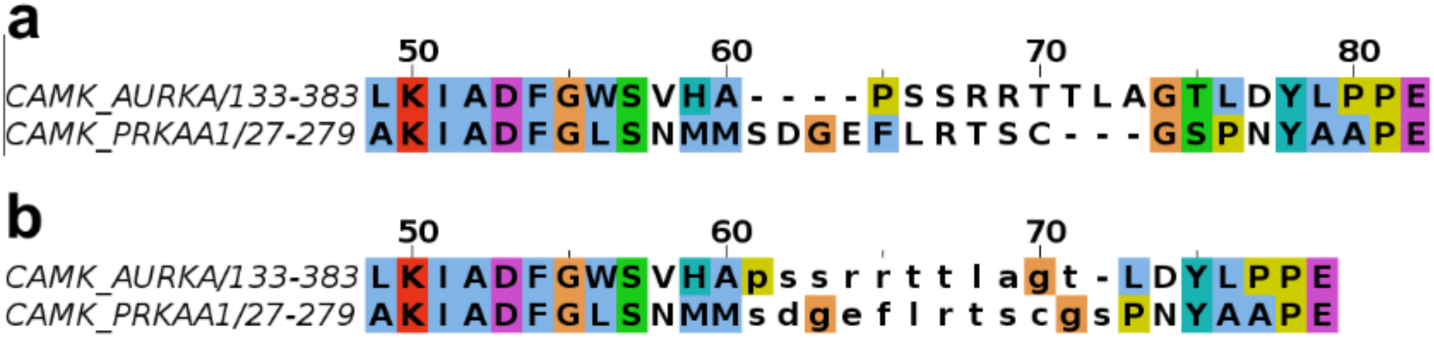
Structural alignment of the activation loop region of CAMK_AURKA and CAMK_PRKAA1 by **a)** FATCAT and **b)** the SE program. Structure alignment programs like FATCAT often introduce gaps in low similarity loop regions to align segments that are not necessarily homologous. The SE program takes coordinates of the superposed structures and produces an optimized sequence alignment with structurally similar regions in upper case and low similarity residues in lower case letters to distinguish them from aligned residues.

**Figure 4:**
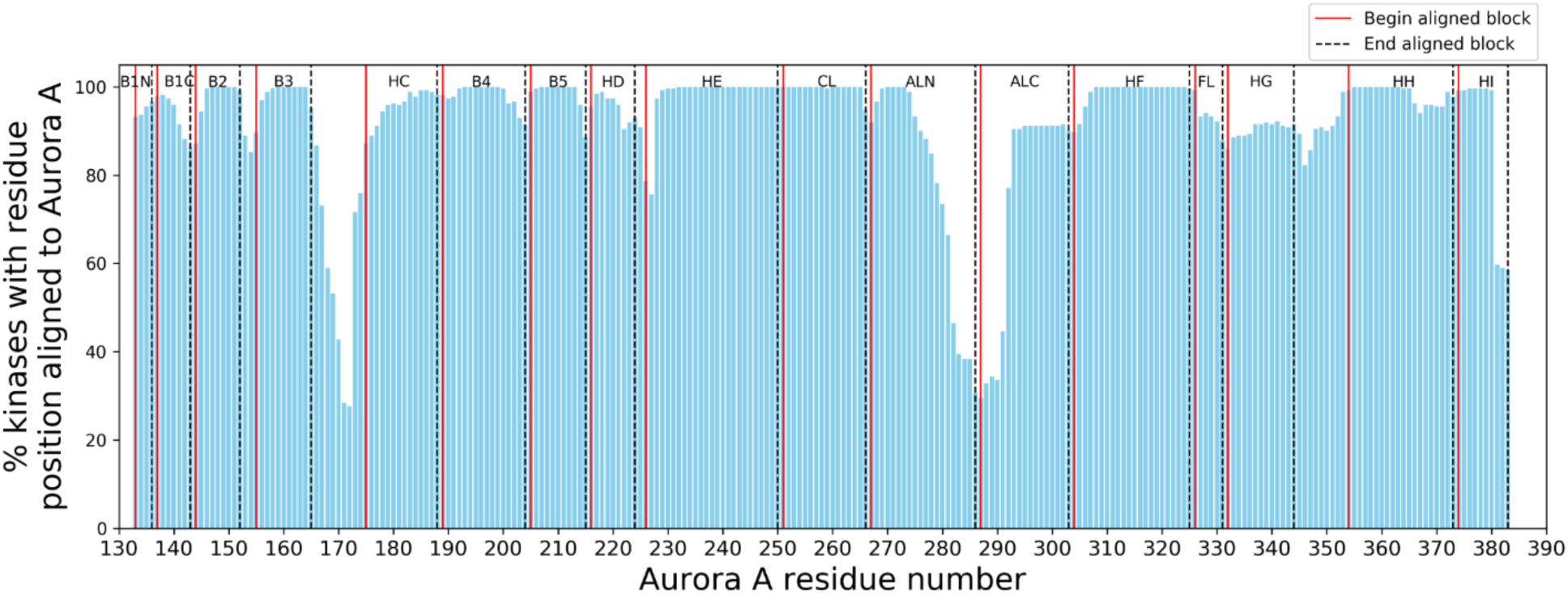
Pairwise structural alignments of Aurora A to 271 human protein kinases using FATCAT. The x-axis represents Aurora A residue numbers; the y-axis displays the number of times each Aurora A residue position is aligned to other kinases in the pairwise alignments. For a description and location of the labeled aligned blocks in Aurora A structure refer to Table 2 and Figure 1. The list of 272 aligned structures is provided in Table S1.

The creation of the MSA was a multi-step process. The initial alignment of all the kinase domain sequences was done using ClustalOmega^37^ which aligned the main conserved regions in a majority of the sequences up to the beginning of the activation loop. Because of very large insertions in the activation loop and in the C-terminal domain in some kinases, the C-terminal domain was aligned only within some families. For example, the AGC-family Great wall kinase (AGC_MASTL) has a 548 amino acid insertion in the activation loop that caused the entire AGC family to be misaligned in the C-terminal domain with respect to the other families.

This alignment was manually edited in Jalview^38^. As shown in Figure 5a, coloring the sequence in “Clustal” format in Jalview greatly aided in adjusting the alignment since it highlighted both level of sequence conservation and physical characteristics of amino acids in each column (hydrophobicity, positive charge, negative charge, etc.). In addition, a file with the secondary structure element boundaries of the kinases of known structure was used to highlight conserved alpha helices and beta sheet strands. We added secondary structure predictions performed with our own secondary structure prediction program based on a deep convolutional neural network and PSI-BLAST and HMM-based sequence profiles. The experimental and predicted secondary structures were visualized in Jalview (Figure 5b). The alignment was improved in cycles of several steps:

**Figure 5:**
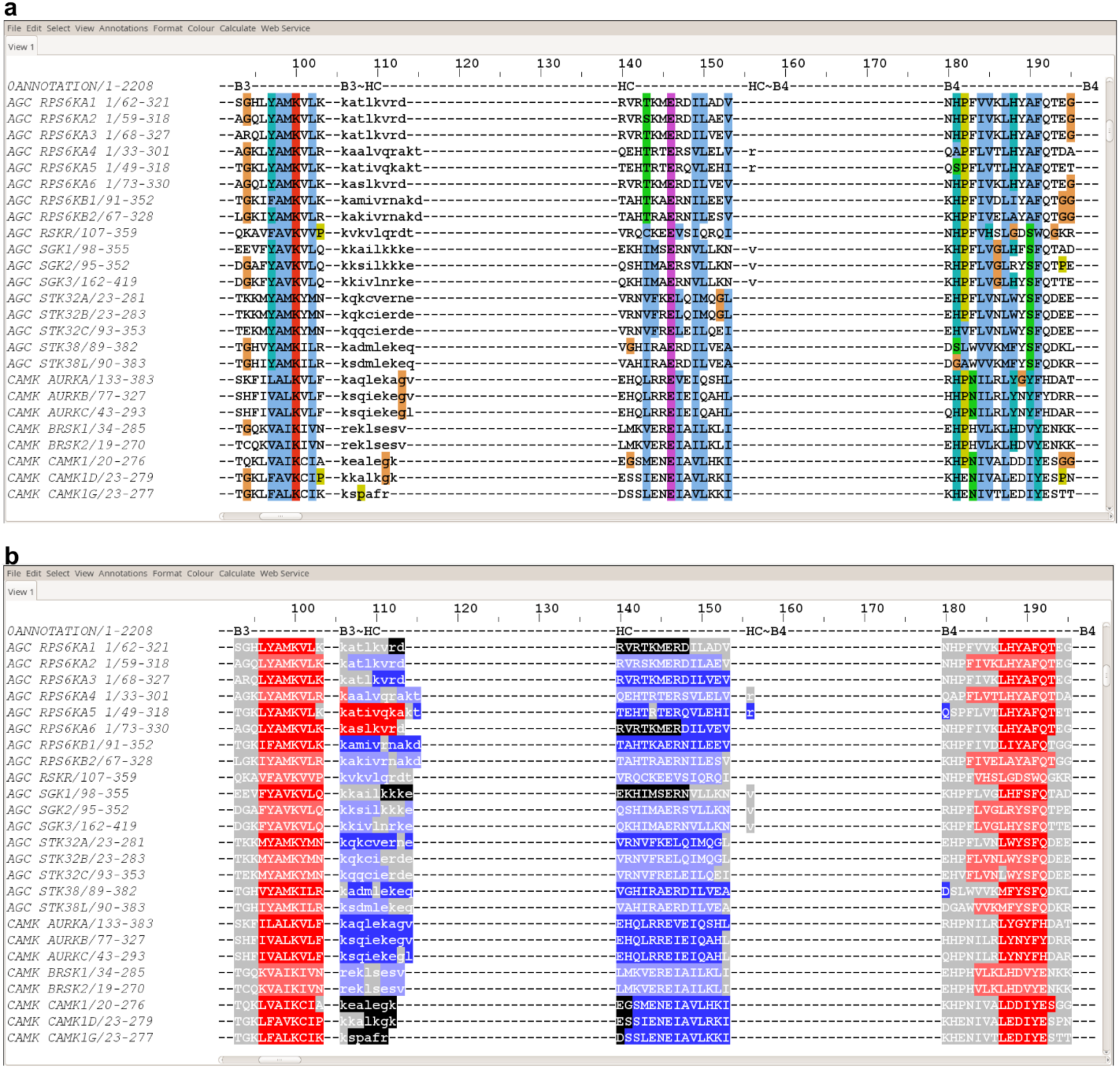
Snapshot of section of N-terminal region of the multiple sequence alignment of human protein kinases displayed using **a)** Clustal coloring scheme in Jalview which colors the residues by their chemical nature if they are conserved or similar to each other; **b)** secondary structure features of proteins. The secondary structure information for proteins with known structures was obtained from PDB files. For the proteins where a crystal structure is not available, secondary structure predictions were performed with our program SecNet (unpublished). The experimentally known and predicted regions are shown in darker and lighter shades of the same color, respectively, with beta sheets in red and helices in blue. Loops are displayed in gray and the residues which were not resolved in crystal structures are colored in black.

1. The main conserved regions identified in Figure 4 were edited to form aligned blocks in the MSA, according to the structure alignments of AURKA N and C terminal domains to the kinases with known structures. These blocks were made as long as possible while respecting the structure alignments.
2. Regions between the aligned blocks that contain variable sequence lengths were left unaligned, and edited in Jalview so that they were left-justified and denoted in lower-case letters. They were separated from the aligned blocks by two empty columns on each end. These blocks were made as short as possible while respecting the structure alignments.
3. The sequences of kinases in each kinase group without known structure were aligned to their closest homologues of known structure, such that all sequence motifs were identified in each kinase. This was greatly facilitated by sorting the alignment by family (AGC, CAMK, etc.) in Jalview, which was possible because of the naming convention described above indicating group membership (AGC_AKT1, etc.) and sorting the alignment by these identifiers.
4. Some kinases without close homologues of known structure required pairwise sequence alignment to other individual kinases with PSI-BLAST and/or hhpred in order to identify regions of the kinase domain that occur after very long insertions or that are highly divergent from other kinases. A small number of kinases were not closely related to any mammalian kinase of known structure, and instead required a sequence alignment to a yeast or insect kinase of known structure, which could then be structurally aligned to AURKA. Examples include: OTHER_TP53RK (33% identity with yeast BUD32, PDB:4WWA^39^); the second kinase domain of OTHER_EIF2AK4 (47% identity to yeast GCN2, PDB:1ZY5^40^); OTHER_PINK1 (47% identity with *Triboleum castaneum* PINK1, PDB:5OAT^41^); and OTHER_PAN3 (49% identical to *Drosophila* PAN3, PDB:4BWP^42^).
5. The G-helix required structure alignments of a region consisting of the G helix itself and the HF∼HG and HG∼HH loops, since the position of the G helix is highly variable within the C-terminal domain of kinases (Figure 1). A small number of kinases do not contain a G-helix as indicated by their experimental structures or those of close homologues (OTHER_BUB1, OTHER_HASPIN) and in a few cases by secondary structure prediction and alignment to more distant homologues (OTHER_RPS6KC1, OTHER_SCYL1, OTHER_SCYL3, OTHER_PEAK1, OTHER_PEAK3, OTHER_PKDCC, OTHER_PRAG1, OTHER_TP53RK, OTHER_PXK, OTHER_RNASEL, OTHER_RPS6KL1, OTHER_POMK, OTHER_STK31, STE_EIF2AK1).

Our final alignment of 497 kinase sequences consists of 2229 columns including all the residue positions and gaps. It has 17 aligned blocks and 16 unaligned low similarity regions. The 17 aligned blocks consist of 8 segments from the N-terminal domain (B1N, B1C, B2, B3, HC, B4, B5, HD) and 9 segments for the C-terminal domain (HE, CL, ALN, ALC, HF, FL, HG, HH, HI). Each block is named for the main element of secondary structure that it contains, although they each contain adjacent loop regions. We name the lower-case insertion regions by the pieces of structure that they connect separated by a tilde, e.g. B2∼B3 is the segment between B2 and B3. B1N and B1C are the N and C terminal segments of beta strand 1; some kinases such as STE_MAP3K8 have an insertion in the strand that produces an extrusion of the chain. CL is the catalytic loop that contains the HRD motif. This segment also contains B6 and B7, which are short beta strands in most kinases. It also contains a 5-residue insertion in one kinase, OTHER_POMK, which is evident in the structure of mouse POMK (PDB:5GZ8^43^). This segment remains part of the aligned region CL, since it occurs in only one kinase. The activation loop is divided into N and C terminal aligned blocks, named ALN and ALC, of 21 and 17 residues respectively. These are long enough to include the common phosphorylation sites of tyrosine kinases (at positions 13 and 14 from the activation loop DFG-Asp) and serine/threonine kinases (at position 12 from the activation loop C-terminus)^44^. Many kinases have long insertions between ALN and ALC. ALN contains the B8 strand prior to the DFG motif at the N-terminus of the activation loop. ALC contains the short EF helix, which extends several residues beyond the APE motif at the C-terminus of the activation loop. FL (“F loop”) is a short beta turn motif with consensus sequence PP[FY] between HF and HG that is conserved in most kinases both in sequence and in structure.

A list of aligned and unaligned blocks is provided in Table 2, including their positions in Aurora A, the column numbers in the MSA, and their length(s). Their positions within the SE alignments are notated in Figure 4. Five of the 16 unaligned regions have nonzero median lengths. The longest of these are the B3∼HC and HG∼HH regions. We left the B3∼HC region unaligned because in most of the 62 AGC and 92 CAMK kinases, the region is in the form of a helix called the B helix while in the other families it takes on a coil form. The HG∼HH unaligned region is highly divergent in structure because of the variation in the position of the G helix. We have created sequence logos with the program WebLogo^45^ to visualize the conservation of residues in all the aligned blocks of our MSA (Figure 6). The logos show the well-known conserved motifs including the HRD motif in the catalytic look and the DFG and APE motifs in the activation loop, as well as hydrophobic positions in the beta sheet strands and alpha helices. For instance, positions 6, 7, and 10 in the G helix contain predominantly hydrophobic amino acids.

**Table 2:**
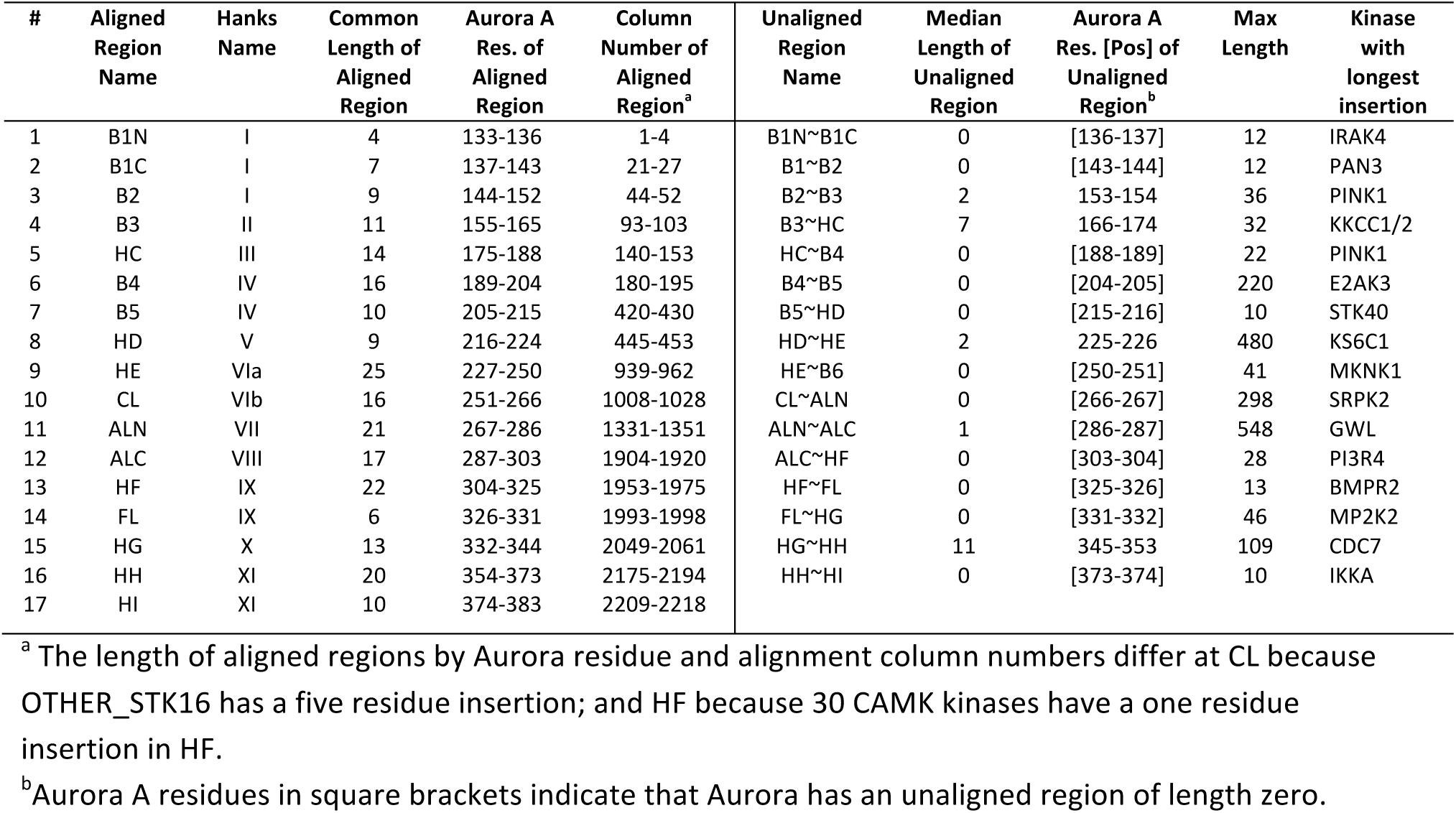
Summary of multiple sequence alignment

**Figure 6:**
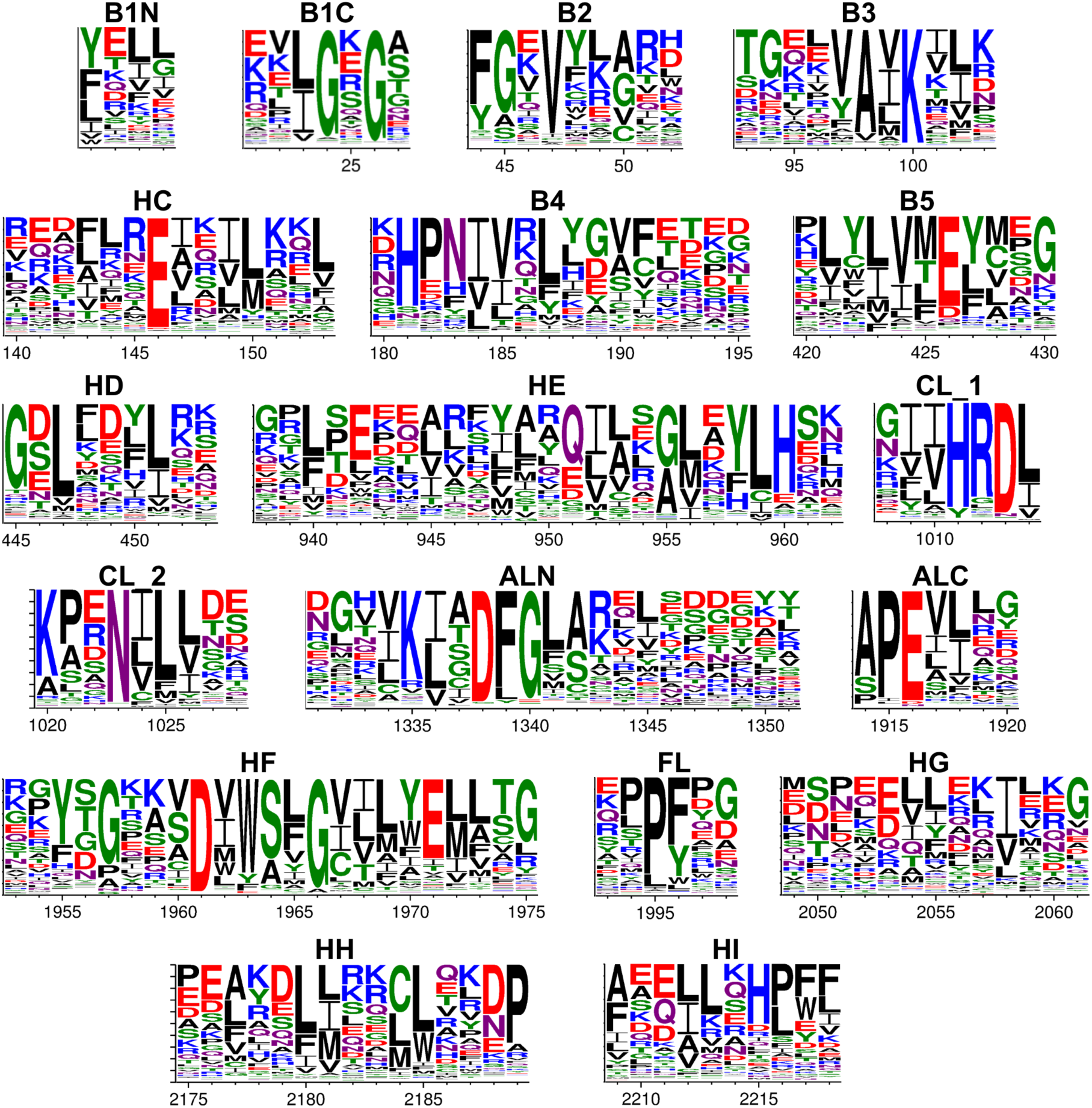
Sequence logos displaying conservation of residues created for all the aligned blocks of the MSA using the webserver WebLogo (http://weblogo.threeplusone.com/). The x-axis represents the column numbers from the MSA. For a description of aligned blocks see Table 2. CL is split into two parts for better visualization.

### Structural validation of the MSA

As described above, the MSA was guided by pairwise alignments of kinase structures to a single kinase (AURKA). However, to determine the accuracy of our MSA we have compared it with the sequence alignments derived from pairwise structure alignments of 272 human kinases in the PDB. Because changes in conformation of the activation loop or movement of the C-helix may affect the corresponding alignment, we used structures that carry an inward disposition of the C-helix as often as possible, as determined by our recent classification of the active and inactive states of kinases^46^. The resulting structure alignments from FATCAT were read by SE to print the unaligned blocks in lower case letters. A residue pair in any two kinases is assumed to be correctly aligned in the MSA if it is also aligned in the pairwise structural alignment of the two kinases.

To perform the validation we have computed three quantities as described in Methods: 1) True positive rate (TPR): the number of residue pairs which are aligned in the MSA and also in pairwise structure alignments divided by the total number of residue pairs aligned in the structure alignments; 2) Positive predictive value (PPV): the number of residue pairs which are aligned in the MSA and also in the structure alignments divided by the total number of residue pairs aligned in the MSA; 3) The Jaccard similarity index^47^: the sum of the number of residue pairs that are identically aligned in both the MSA and the structure alignments divided by the total number of unique residue pairs aligned in the MSA or the structure alignments or both (counting each pair only once). The Jaccard index shows the overlap between MSA and structural alignments, and penalizes both under- and over-prediction of aligned residues in the MSA.

The average values and distributions of these quantities are presented in Table 3 for our MSA. We have also compared the quality of our alignment with the previously published alignments by Manning et al.^5^, Möbitz^16^, Kwon et al.^23^, a hidden Markov model (HMM) derived from our MSA, and the initial ClustalOmega alignment. The average TPR for our MSA is 0.97, which is significantly better than the Möbitz (0.88), Kwon (0.90), and Manning (0.80) MSAs. The initial ClustalOmega alignment had a TPR of only 0.74.. Similarly, our MSA also has the highest PPV (0.97) and the highest value for the Jaccard index of 0.94. The HMM derived from our MSA has TPR of 0.92, which is less accurate than the MSA itself but more accurate than the Möbitz, Manning, and Kwon alignments. It may therefore be of use in aligning kinases from other species to our MSA of human kinases.

**Table 3.** Average values for TPR, PPV and Jaccard similarity

|  | Kinases in alignment | Kinases in validation | TPR | PPV | Jaccard similarity | Gap regions per kinase |
| --- | --- | --- | --- | --- | --- | --- |
| **FoxChase\*** | 497 | 272 | 0.969 | 0.967 | 0.938 | 18.7 |
| **HMM\*\*** | 497 | 272 | 0.922 | 0.919 | 0.857 | - |
| **Manning** | 491 | 272 | 0.799 | 0.809 | 0.694 | 44.6 |
| **Möbitz** | 489 | 271 | 0.882 | 0.880 | 0.799 | 19.9 |
| **Kwon** | 494 | 271 | 0.901 | 0.908 | 0.835 | 139.6 |
| **Clustal** | 497 | 272 | 0.739 | 0.754 | 0.612 | 22.3 |

\* Our alignment
\*\* HMM derived from our alignment
TPR = true positive rate
PPV = positive predictive value

We calculated the ‘gappiness’ of each element, which we identify as the average number of gap regions in each sequence in the MSA. These are also contained in Table 3. Our alignment is the least gappy, with average number of gap regions of 19. While we have 16 unaligned regions, three of the aligned regions contain short gapped regions internally to accommodate one or more kinases with an unusual insertion in the aligned region. The Möbitz and ClustalOmega alignments are slightly more gappy than ours, while the Manning and Kwon alignment are substantially gappier with 45 and 140 gap regions per sequence respectively.

We examined the positions where the structure alignments and our MSA are discrepant (Figure 7a). In some cases, the discrepancies occur in positions within the aligned blocks that are immediately adjacent to the unaligned segments. This is because the positions of the aligned blocks are not ideal for every single kinase but are a form of consensus position. For example, 30% of the alignments have a discrepancy near the B4/B5 boundary. In the remaining cases, they represent structural shifts of elements of secondary structure in some kinases relative to the kinase domain. This is particularly true of the C-helix, which shows a discrepancy between the sequence alignment and the structural alignment at a rate of about 5%. Examination of cases where the C-helix is misaligned indicates that the sequence alignments are probably correct, and the differences with the structure alignment are because of shifts in the position or orientation of the C-helix relative to the rest of the N-terminal domain. For example, the C-helix is misaligned in 50% of the structure alignments of MAP2K1 and MAP2K2, despite the fact that the conserved C-helix glutamic acid that forms a salt bridge with a lysine residue in the B3 strand in most kinases is correctly aligned in our MSA. Another example is the C-helix of STK38 (PDB: 6BXI^48^), which is rotated by 100°, and thus the conserved glutamic acid residue and the entire C-helix do not align with the homologous residues in closely related kinases. An active form structure of STK38 is not available, but is predicted to have the C-helix salt bridge to the lysine residue in the β3 strand^48^.

**Figure 7:**
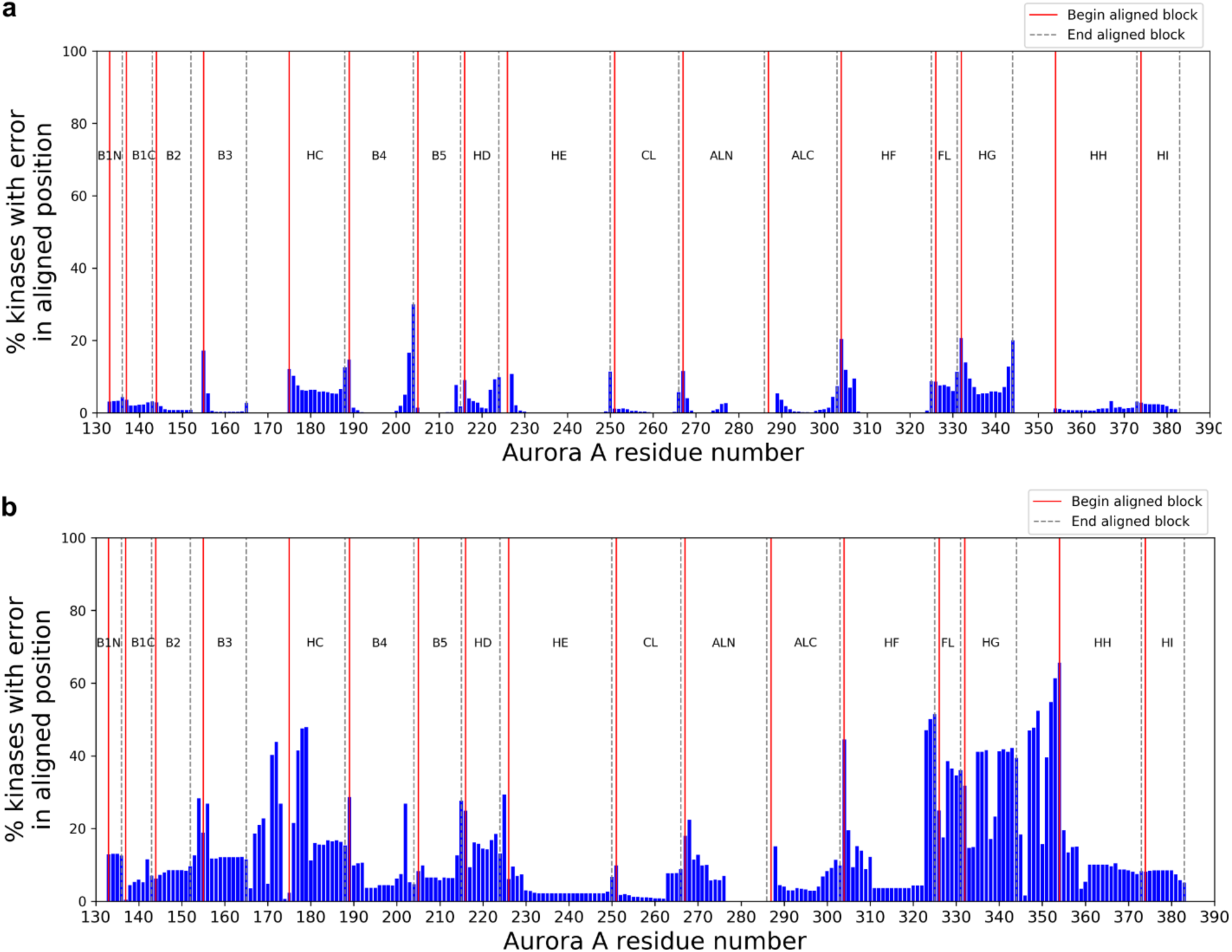
Histogram displaying the discrepancy between benchmark pairwise structural alignments of kinases and **a)** FoxChase and **b)** Manning’s multiple sequence alignment. The y-axis represents the number of pairwise alignments when a residue pair is aligned in the benchmark but is not aligned in the MSA. The values are plotted with Aurora A residue numbering as a reference on the x-axis.

We performed the same analysis of the alignment errors for the Manning et al. alignment (Figure 7b). While there are more misalignments throughout the kinase domain in the Manning alignment than in ours, the G-helix region is misaligned in about 50% of the pairwise alignments in comparison to the structure alignments. This is likely due to lower sequence conservation in this region and the absence of readily identified sequence motif, like the HRD and DFG motifs. The segment before the C-helix is also poorly aligned compared to structure alignments because of the presence of the B helix in AGC and CAMK kinases and coil region elsewhere, as noted above.

Some kinases have very long insertions that have confounded some previous multiple sequence alignments. AGC_MASTL (Great wall kinase) contains a 548-residue long insertion within the activation loop (residues 188-735) and is the longest kinase domain sequence at 801 amino acids. In the Manning and Möbitz alignments, the kinase domain is defined as residues 35-310, and both miss the entirety of the C-terminal domain that follows the activation loop. OTHER_RPS6KC1 (Ribosomal protein S6 kinase delta-1) has a 480-residue long insertion (residues 419-898) in the HD∼HE unaligned region. In the Manning alignment, the kinase domain begins on residue 822; in our alignment it begins at residue 337. OTHER_CDC7 has a 98-residue long insertion (residues 441-539) between HG and HH, while the kinase domain is defined as residues 58-472 in the Kwon alignment, thus missing the HH and HI helices.

### Phylogenetic trees and group membership

In their paper on the human kinome, Manning et. al provided a phylogenetic tree and classified the human protein kinases into nine groups extending the early Hanks^49^ and Hunter^50^ schemes. These groups consisted of AGC, CAMK, CK1, CMGC, NEK, RGC, STE, TKL and TYR. A total of 83 protein kinases were placed in OTHER category because no significant relationship to any of the nine groups was recognized. However, this classification was done with only a limited amount of structural information, and as shown above, the Manning multiple sequence alignment was only 80% correct. We have revisited the phylogeny and classification of kinases to see if we can assign groups to some of the OTHER kinases by benefiting from our structure guided multiple sequence alignment.

Because using aligned blocks tends to result in better phylogenies^51^, we built the tree with the 17 conserved blocks from the alignment (Figure 8). The tree was created using the neighbor joining algorithm in the software MegaX^52^ and was visualized using the webserver iTOL^53^. To test the robustness of the tree we computed branch supports using a bootstrap calculation by the program Booster^54^. It uses a gradual expectation function to quantify the presence of a branch in replicate trees. The bootstrap value represents the percent of replicate trees in which a specific branch order was observed. A branch with a bootstrap value of 70% or above is considered robust and representative of the information in the sequence alignment. We have observed that in our phylogenetic tree most of the internal branches have a value of 70 or above. The resulting tree clusters most of the kinases into the previously recognized nine groups. Uniprot includes the NEK kinases as a separate group, which also appears in our tree. In our tree, the RGC kinases form a small sub-branch within the TKL group, but we have retained the designation.

**Figure 8:**
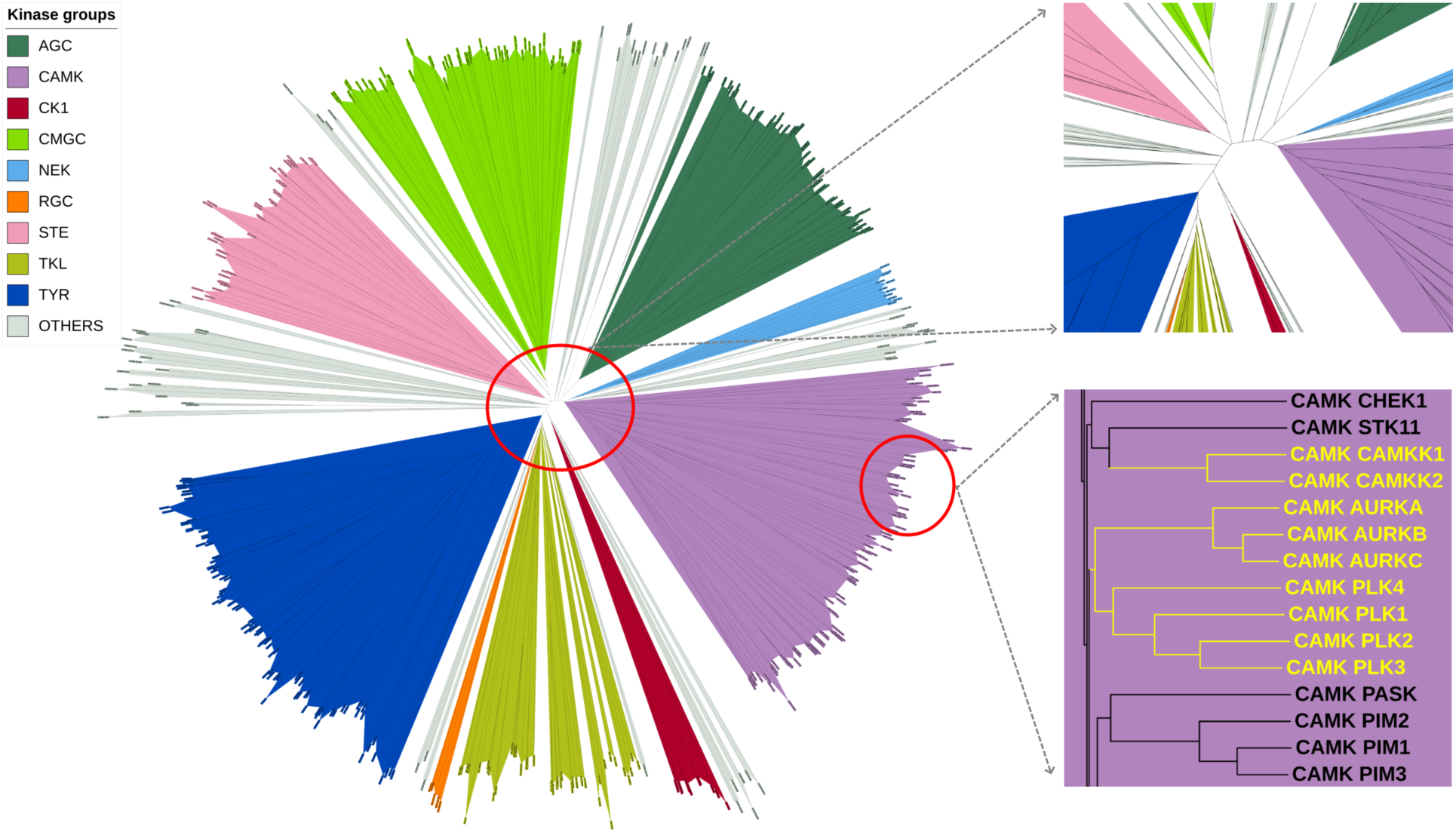
Unrooted phylogenetic tree of human protein kinases created by using our MSA. The nine kinase groups are displayed in different colors with OTHER in gray. The center of the tree is magnified on the right. Nine of the ten OTHER kinases assigned to CAMK group by our analysis are shown as a dendrogram on the right side (PLK5 is a truncated kinase domain and is not shown in the tree). The figure was created using the iTOL webserver and can be accessed at https://itol.embl.de/shared/foxchase

Among the kinases which are assigned to a group by Uniprot, we observed that eight STE kinases, MAP3K7 (TAK1 in Manning), MAP3K9 (MLK1), MAP3K10 (MLK2), MAP3K11 (MLK3), MAP3K12 (ZPK), MAP3K13 (LZK), MAP3K20 (ZAK), and MAP3K21 (MLK4) form a tight cluster in the TKL group branch. These were also in the TKL branch of the Manning tree. Similarly, six sequences consisting of the second domains of RPS6KA1, RPS6KA2, RPS6KA3, RPS6KA4, RPS6KA5, and RPS6KA6 which were annotated to be in the AGC group by Uniprot also cluster in CAMKs (as they are in the Manning tree). The first domains of these kinases are AGC members.

Because of their remote homology, most of the OTHER kinases branch out into separate clades in between the major groups. Due to the smaller size of these clades and relatively low similarity between the members we have not classified them as individual kinase groups. In our tree there are seven OTHER kinases in Uniprot that are correctly assigned to groups by Manning. Four kinases -- STK32A, STK32B, STK32C, and RSKR, form a branch within the AGC group. These kinases were also classified as AGCs by Manning et. al. (labeled YANK1, YANK2, YANK3, SgK494 respectively). Three kinases, CSNK2A1, CSNK2A2, and CSNK2A3, form a tight cluster within the CMGC group. These are listed as OTHER by Uniprot and while CSNK2A1 and CSNK2A2 were designated CMGCs by Manning. One kinase, PBK (also called TOPK) is assigned to STE in Uniprot, but to OTHER by Manning. In our tree, it sits just outside the TKL group but we would still classify it as OTHER.

We have identified a set of ten kinases from the OTHER category in both Manning and Uniprot that can be appropriately assigned to the CAMK group. They form a branch in the middle of the CAMK group by nine kinases consisting of AURKA, AURKB, AURKC, CAMKK1, CAMKK2, PLK1, PLK2, PLK3, and PLK4 (Figure 8). PLK5 is a pseudokinase consisting only of the C-terminal domain, although mouse PLK5 is full-length^55^. We have included it in the CAMK group because of its close sequence relationship with the other PLKs.

To confirm the changes in group membership, we created HMM profiles for each of the nine groups of kinases as defined by Manning et al. We then scanned each of the 497 kinase sequences against the nine group HMM profiles. A cutoff score of 200 was consistent with the assignments by Manning except for the changes described above. The novel assignments are the ten kinases that we can confidently move from OTHER to CAMK described above. The HMM scores clearly assign them to CAMK rather than AGC or OTHER, since the new CAMKs cluster with the other CAMK kinases (Figure 9).

**Figure 9:**
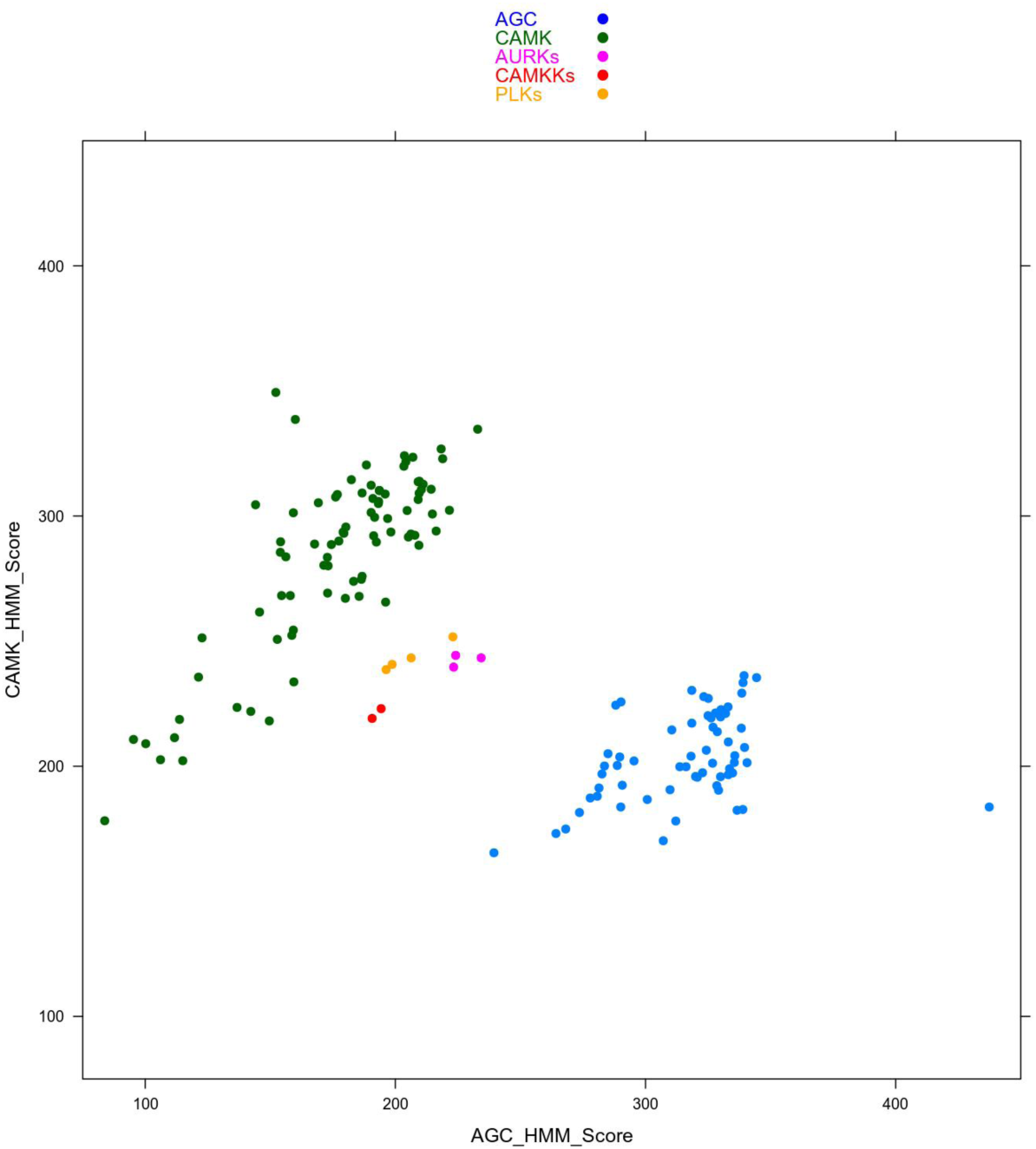
HMM scores of CAMK and AGC kinases. HMMs were built on our alignment of AGC and CAMK kinases according to the original Manning assignments. Three subfamilies of OTHER kinases have CAMK scores in the same range as other bona fide CAMK kinases (green). These scores are higher than their scores with the AGC HMM (blue). These kinase subfamilies consist of Aurora (AURKA, AURKB, AURKC), CAMKK (CAMKK1 and CAMKK2), and PLK (PLK1, PLK2, PLK3, PLK4).

Using the group assignments after correcting Uniprot, and new assignments from our analysis in the AGC, CAMK, CMGC and TKL groups, we created new HMM profiles to identify if any OTHER sequences could also be reassigned. However, in the second iteration none of the OTHER category kinases exhibited high scores against any group HMM profile.

## DISCUSSION

Typical kinase domains possess a well-defined fold that is similar across the available structures. Many kinases are not well studied – the so-called dark kinome^56^, and it is possible to generate hypotheses about their sequence-structure-function relationships by examining their phylogenetic and structural relationships to well-studied kinases. To enable this and for many other purposes, we have created a structurally-validated, multiple sequence alignment of 497 human protein kinase domains—fully annotated with gene, protein, group name, UniProt accession identifiers, and residue numbers. The MSA contains 17 aligned blocks of conserved elements of typical kinase domain and 16 intermittent low similarity regions with varying length insertions. Our aim was to create a parsimonious alignment without unnecessary gaps; the residues in low similarity regions were therefore not aligned but formatted as left-justified blocks of lower case letters to distinguish them from aligned regions. It is reminiscent of the first multiple sequence alignment of kinases produced by Hanks et al. in 1988^49^, which the authors also described as “parsimonious.”

Alignments are only useful if they are accurate. While several multiple sequence alignments of human kinases have been published and are available online^5,16,19,23^, none of them has been structurally validated. We assessed the accuracy of our alignment with a set of all-against-all pairwise structural alignments of 272 human kinases, and calculated true positive rates (TPR), positive predictive values (PPV), and the Jaccard similarity index. In a large-scale benchmark of sequence alignment methods, we referred to them as *f*_*D*_ (for developer) and *f*_*M*_ (for modeler) for TPR and PPV respectively^57^. Yona and Levitt subsequently used these values (renamed *Q*_*D*_ and *Q*_*M*_) to benchmark profile-profile sequence alignments, and added *Q*_*C*_ or *Q*_*Combined*_^58^, which is simply the Jaccard index.. The Jaccard index penalizes both overprediction and underprediction in our sequence alignments. We used all three values for our alignment (0.97, 0.97, and 0.94 respectively) to demonstrate that our alignment is more accurate than the others available.

The errors in our MSA of kinases are mostly limited to the boundaries of conserved blocks where the variability of residue positions across kinases make their unambiguous placement in aligned blocks difficult. However, structure alignments do not always align every homologous pair of residues in two proteins. This occurs when residues are disordered in one of the structures or where there is significant conformational change. In a small number of kinases the only structure available has a significantly rotated C-helix. Structure alignment therefore sometimes does not align the homologous residues of the C helix in two kinases. The same is true for the G-helix in some kinases, which may be positioned in different locations within the C-terminal domain, but retains a homologous sequence and structure, and thus is aligned differently in our MSA than in the structure alignments.

While our MSA is guided by predicted secondary structure of kinases and benchmarked with pairwise structural alignments, there are other approaches like ConTest and QuanTest which conversely use prediction of contact maps and secondary structure to assess the quality of the alignment^59,60^. The predictions which are close to the values from experimental structure suggest that the multiple sequence alignment is more accurate.

With a more accurate MSA in hand, we have revisited the phylogenetic tree of human kinases. The widely used phylogenetic tree of Manning et al. is based on an alignment that has TPR and PPV values of only 0.80 and 0.81. Uniprot also provides a classification of kinases into the same groups as Manning et al (with the addition of NEK kinases). The Uniprot annotations that differ from Manning et al. are all incorrect. Fourteen kinases are placed in the wrong groups by Uniprot, and another seven are placed in OTHER but can easily be placed within one of the defined groups, in agreement with the Manning annotations. One kinase (PBK or TOPK) is labeled as STE by Uniprot but we agree with Manning’s annotation of OTHER.

Of greater interest, our phylogenetic tree and hidden Markov models for each group can help us assign ten kinases to the CAMK group; these are listed as OTHER by Manning et al. and Uniprot: AURKA, AURKB, AURKC, CAMKK1, CAMKK2, PLK1, PLK2, PLK3, PLK4, and PLK5. Manning et al. placed the CAMKK and PLK kinases adjacent to the CAMK group and the Aurora kinases at the base of the AGC branch (but did not designate them as AGC). From our hidden Markov model of CAMK kinases and a phylogenetic tree based on our MSA, these kinases fit clearly into the CAMK group. Experimental data confirm that these assignments are correct. CAMKK1 and CAMKK2 (Ca^2+^/calmodulin-dependent kinase kinase 1 and 2) both phosphorylate Ca^2+^/calmodulin-dependent kinases and bind calmodulin^61-63^. There is also direct evidence of calmodulin binding to PLK1^64^ and of a calmodulin homologue, calcium-and-integrin-binding protein (CIB), to both PLK2 (Snk in the kinome poster) and PLK3 (Fnk)^65^.

Manning et al. put the Aurora kinases on the AGC branch (but not labeled AGC), which is closely related to the CAMK group. Both groups possess a B helix that is not present in the other families. The HMM and the phylogenetic tree show that the three Aurora kinases fit better into the CAMK group than the AGC group. In an earlier study with colleagues, we have shown experimental evidence that Aurora A binds calmodulin^66^, supporting its assignment to the CAMK group. Calmodulin also binds to Aurora B kinase (AURKB), preventing its degradation via the E3 ligase FBXL2 subunit^67^.

Our MSA provides the benefit of a common numbering scheme using the columns of the alignment facilitating comparison across all the kinase sequences. The identification of equivalent residue positions helps in generalizing experimental data from one kinase to another. For example, substrate specificity is highly correlated with the amino acid type at a small number of positions within the substrate binding site^68^. Creixell found that specificity could also be modulated by more remote sites^19^, based on a multiple sequence alignment of kinases derived with ClustalOmega^37^. It is likely that our more accurate alignment would facilitate this analysis and produce more reliable predictions. Other areas where an accurate alignment and phylogeny may be useful are in predicting inhibitor specificity^69^, regulatory mechanisms through protein-protein interactions, and computational protein design of kinases with altered functionality^70^.

Our alignment is included as supplemental data and on our website, and will be updated as new structures are determined. We hope that it will be of use in kinase biology and therapeutic development.

## METHODS

### Identification of human protein kinases

The list of human typical and atypical protein kinases was obtained from Uniprot website (https://www.uniprot.org/docs/pkinfam). To identify any unlisted kinases we searched human sequences in Uniprot with PSI-BLAST using the typical and atypical protein kinase on the Uniprot page as queries. PSI-BLAST was also used to identify structures of human kinases or their closest homologues in the PDB. The structures of atypical kinases (or homologues thereof) were examined structurally using a stand-alone version of FATCAT provided by Adam Godzik (personal communication) and structural superposition by CEalign in Pymol to Aurora A. Four of the atypical kinase families are visibly related to typical kinases, but contain significant fold differences. The other two families are not homologous to typical kinases.

A total of 497 typical kinase domain sequences from 484 kinase genes (13 genes have two kinase domains each) were used to create the MSA. These sequences were initially divided into 9 phylogenetic groups as per the Uniprot nomenclature: AGC, CAMK, CK1, CMGC, NEK, RGC, TKL, TYR, and STE, and a tenth group of diverse kinases designated OTHER. Gene names were retrieved from the Human Gene Nomenclature Committee website (http://genenames.org)^35^. Each kinase sequence was labeled by group name underscore HGNC gene name, for example AGC_PRKACA for KAPCA_HUMAN. The 13 kinases that have two kinase domains in the polypeptide chain were labeled with an underscore, for example TYR_JAK1_1 and TYR_JAK1_2. The boundaries were determined with PSI-BLAST of the full-length Uniprot sequence against the PDB.

### Multiple Sequence Alignment

The kinase sequences (except some with very long insertions like GWL) were aligned using ClustalOmega^37^ to prepare an initial alignment. This was manually edited using Jalview^38^ to make sure that conserved motifs such as the DFG and HRD motif were aligned across most of the sequences. The sequences with low sequence similarity to most of the other kinases and those containing long insertions were difficult to align. To improve the accuracy of the alignment, pairwise structural alignment of the kinases which have a crystal structure was performed using the structure of Aurora kinase (3E5A_A) and the program FATCAT^71^. However, for the kinases where a structure was not known, alignment of the closest known structures to Aurora A were used to edit the alignment with Jalview In a few cases, the most closely related structures were not human or even mammalian kinases. In these cases, the non-human kinase was structurally aligned to Aurora A and the target kinase was added to the MSA by transitive alignment. For a few distant kinases where a closely related structure or sequence was not known, HHPred was used to identify similarity to another kinase^72^.

### Structural validation of the MSA

The MSA was structurally validated using a set of pairwise structural alignments as a benchmark. The benchmark consists of all vs all pairwise structural alignments of 272 kinases with known structures in the PDB. Two kinases were excluded: the BUB1B kinase domain in the cryo-EM structure of the anaphase promoting complex (PDB: 5KHU, chain Q^73^) is completely disordered; the structure of PBK (TOPK_HUMAN, PDB: 5J0A^74^) contains two monomers with half of the N-terminal domains of each chain swapped with the other monomer, making it difficult to align to other kinase domain structures. The structure-based sequence alignments were created using FATCAT in rigid mode and optimized using SE^36^. For kinases with multiple structures known the structure for validation was selected based on their conformational states using our previously published nomenclature^46^. The active state BLAminus conformation was preferred over others, followed by different kinds of DFGin inactive states - ABAminus, BLAplus, BLBminus, BLBplus, BLBtrans and DFGout-BBAminus.

A residue pair between two kinases in the MSA was considered to be aligned if it was also aligned in the benchmark pairwise structural alignment of the same kinases. Using this information, the accuracy of the MSA was assessed by computing three quantities TPR, PPV and the Jaccard similarity index. For each pair of sequences, we first calculate the number of aligned residue pairs that are present in both the sequence alignment and the structure alignment (*N*_*correct*_). The TPR is the ratio of *N*_*correct*_ and the number of residue pairs aligned in the structure alignment (*N*_*struct*_). For computation of the TPR, residue pairs in the structure alignment are skipped if either or both residues are contained in the unaligned (lower-case) blocks of the sequence alignment. This takes care of situations that occur when two kinases have identical length segments between two of our aligned blocks; the structure alignment program would align them but they would be indicated as unaligned in our sequence alignment. The alignment of Kwon et al. also includes unaligned regions in lowercase and is treated in the same way.

PPV is the ratio of *N*_*correct*_ and the number of aligned residue pairs in the sequence alignment (*N*_*seq*_). For the PPV, residue pairs in the aligned blocks of the sequence alignment are skipped if one or both residues are aligned to gap characters in the structure alignment. This is usually either because the residues are disordered (no coordinates) in one of the structures or because there is a significant conformational change of a loop and the residues are aligned to gaps. The Jaccard similarity index is the ratio of *N*_*correct*_ and the number of unique aligned pairs in either the structure alignment or the sequence alignment (counting each only once). For the Jaccard index, all the pairs skipped in TPR and PPV are also skipped. A script for calculating these values is available on https://github.com/DunbrackLab/Kinases. The aligned residues from the pairwise structure alignments are available on https://zenodo.org/record/3445533 (DOI 10.5281/zenodo.3445533).

We also compared our MSA accuracy with the previously published alignments. These alignments did not contain residue ranges in the Uniprot sequences, and used different nomenclature for the protein names. To identify a correspondence between the sequences in previously published alignments and our MSA, we performed PSI-BLAST searches of each of their sequences against Uniprot and renamed them according to our scheme (groupname_genename).

### Phylogenetic tree

The phylogenetic tree of human protein kinases was created from an MSA obtained by deleting the unaligned regions from our MSA. A distance matrix using the p-distance in the program MegaX^52^ was created which was used to create a phylogenetic tree with the neighbor-joining algorithm.

To perform bootstrap analysis on the phylogenetic tree we generated 5000 bootstrap alignments using the program goalign (https://github.com/evolbioinfo/goalign). The alignments were read by MegaX using the same algorithm as mentioned above to infer bootstrap trees. Finally, ‘transfer bootstrap’ values were computed using the stand-alone version of the program Booster (https://booster.pasteur.fr/)^54^.

The tree was saved in Newick format and uploaded to iTOL webserver for visualization where each clade was colored according to its kinase group (https://itol.embl.de/shared/foxchase). It can be visualized in rooted and unrooted representations with bootstrap values using the buttons provided by iTOL interface.

### HMM profiles of kinase groups

We used the hmmbuild program of the HMMER3 package^75^ to create HMM profiles for each of the nine kinase groups as defined by Manning et al. The input MSA for each group was extracted from the main MSA and any empty columns were deleted. All 497 kinase sequences were run against the nine HMM profiles using the program hmmsearch^75^. The scores of each sequence against nine HMMs were sorted and the group with highest score against each sequence was identified.

## Supporting information

Supplemental Table S1.txt

Human Protein Kinase Domain sequences - fasta format

Multiple Sequence Alignment - Human Protein Kinases - fasta format

Multiple Sequence Alignment - Human Protein Kinases - Clustal format

## Acknowledgments

This work was supported by NIH grant R35 GM122517 to RLD. We thank Maxim Shapovalov for the program *SecNet* for prediction of protein secondary structure.

## Supplemental Files

TableS1.txt – data on each of 497 kinases in the alignment in text format

TableS1.xlsx -- data on each of 497 kinases in the alignment in Excel format

Human-PK-v1.aln – our alignment in Clustal format (version 1)

Human-PK-alignment-v1.mfa – our alignment in FASTA format, suitable for visualization in Jalview

Human-PK-sequences-v1.fasta – the domain sequences of 497 kinases (unaligned)

